# SplineDist: Automated Cell Segmentation With Spline Curves

**DOI:** 10.1101/2020.10.27.357640

**Authors:** Soham Mandal, Virginie Uhlmann

## Abstract

We present SplineDist, an instance segmentation convolutional neural network for bioimages extending the popular StarDist method. While StarDist describes objects as star-convex polygons, SplineDist uses a more flexible and general representation by modelling objects as planar parametric spline curves. Based on a new loss formulation that exploits the properties of spline constructions, we can incorporate our new object model in StarDist’s architecture with minimal changes. We demonstrate in synthetic and real images that SplineDist produces segmentation outlines of equal quality than StarDist with smaller network size and accurately captures non-star-convex objects that cannot be segmented with StarDist.

## 1. INTRODUCTION

A vast amount of biological questions investigated using microscopy data rely on an initial partitioning of the image into individual objects, a task referred to as instance segmentation [1]. Dividing an image into object regions multiplies the amount of data points that can be extracted from it. Protein expression [2] and morphology [3], among many others, can then be quantified at the single-cell level, thereby moving away from population averaging. Due to the sheer amount of imaging data acquired in modern biology experiments and because segmentation is so essential, the development of automated computational segmentation methods has been at the heart of bioimage analysis since its early days [4]. In the past decade, deep learning approaches have revolutionized this field of research [5]. Convolutional neural networks (CNN) in general, and the U-net model in particular [6], have demonstrated an unprecedented capability to consistently produce excellent segmentation results in a wide range of data, massively reducing the need for manual intervention.

Recently, StarDist [7] established itself as a popular detection and instance segmentation tool in the bioimaging community due to its simplicity and efficiency. StarDist generates object outlines as star-convex polygons: it relies on a U-net architecture to predict the locations of a fixed number of points on an object’s contour, and constructs the final outline by linearly interpolating between them. This object representation is reminiscent of the one used in the parametric B-spline active contour (AC) algorithm [8] and in its many variants [9, 10, 11]. In spline AC methods, objects are segmented with a spline interpolated curve that deforms in the image following the minimization of a handcrafted energy functional. Beyond segmentation itself, the main advantage of these approaches is to provide a continuously-defined description of the object contour as a parametric curve that can readily be used to, *e.g.*, perform statistical shape analysis [12]. The flexibility of spline curves makes them suitable for a variety of segmentation problems, but their optimization requires domain-expertise, restricting their use in practice. In preliminary work, we proposed a first step towards circumventing this limitation by directly predicting spline interpolated outlines from an input image using a CNN regressor [13]. Our prototype was however limited to binary images featuring a single object.

Here, we introduce SplineDist, an instance segmentation algorithm that combines the capabilities of StarDist with those of spline approximation. Thus, SplineDist benefits both from the detection performance of StarDist and the segmentation quality of spline models. While [13] could only produce spline outlines from binary masks of single objects, SplineDist is not bounded by any such restriction. It can handle raw images featuring many objects and therefore provides end-to-end instance segmentation. We show that SplineDist equals StarDist’s excellent segmentation results with a smaller number of parameters. Besides, SplineDist can segment objects regardless of their star-convexity, thus lifting StarDist’s main restriction. As a result, SplineDist expands the range of applicability of StarDist and offers a robust alternative to spline-based AC methods.

The paper is organized as follows. In Section 2, we recall the technical details of StarDist and describe the construction of SplineDist. We then experimentally compare the two methods and illustrate SplineDist’s capabilities in Section 3. Finally, we conclude the paper in Section 4.

## 2. METHODS

### 2.1. StarDist

Given an input image featuring one or many biological objects, StarDist [7] predicts a star-convex polygon of *R* vertices for every pixel. More precisely, at every pixel location (*i,j*) in the image, StarDist predicts the set of radial distances 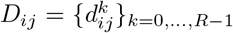 to the boundary of the object enclosing (*i,j*) at *R* equidistant angles 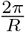 (Figure 1b). Because the distances 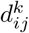 are not defined for background elements, StarDist simultaneously predicts an object probability *p_ij_* for each pixel, allowing to ignore proposals from pixels with low object probability. For each pixel in the image, StarDist’s loss is formulated as

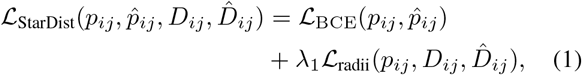

where *p_ij_* and 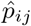 are the ground truth and predicted pixelwise object probabilities, and *D_ij_* and 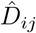 are the ground truth and predicted sets of radial distances to the object contour. The first term, 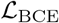, is the standard binary cross entropy loss [14] and λ_1_ is a regularization factor. The second term, 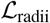, is a mean absolute error loss weighted by ground truth object probabilities and expressed as

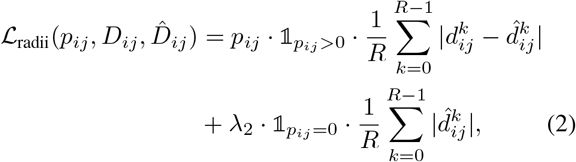

with λ_2_ a regularization factor. This loss is finally averaged over all the pixels present in the image. An important consequence of (2) is that, during training, a set of individual ground truth radial distances *D_ij_* must be generated for each individual pixel (*i,j*) from a reference instance labelling of the entire image.

**Fig. 1.**
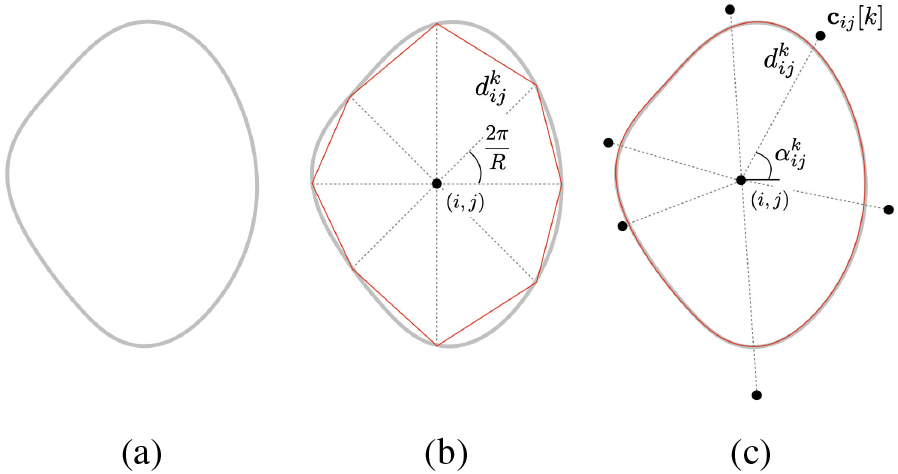
StarDist and SplineDist object models. (a) A reference object outline. (b) For each point (*i, j*) enclosed in the object, StarDist models the outline as a star-convex polygon of *R* points located at fixed, uniformly distributed angles and radial distances 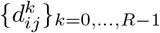. (c) In contrast, SplineDist models the outline as a planar spline curve (3) generated by *M* control points 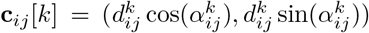. Here, *R* = 8, *M* = 6, and *φ* = *β*^3^ the cubic B-spline basis.

Once star-convex polygons have been predicted for all the pixels, overlapping candidates are filtered by a non-maximum suppression (NMS) step taking into account their associated object probabilities. The final set of polygons corresponds to individual object instances in the image.

### 2.2. SplineDist

Instead of representing objects as polygons, we propose to adopt the more flexible model used in spline AC algorithms, which uses spline interpolation. Essentially, SplineDist models an object outline as a two-dimensional parametric planar spline curve s defined by

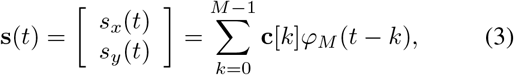

where 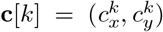 are two-dimensional control points, which are the parameters of the spline model. While the number *M* of control points is fixed, they can be located anywhere in the image. The function 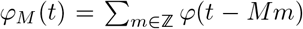 is the *M*-periodized version of a predefined spline basis φ, allowing to get a closed curve [9]. This object model is general and unifying: we can indeed consider any spline basis for φ, typically polynomial B-splines of any order. In the following, we denote as SplineDist _*β^n^*_ the version of SplineDist relying on *φ* = *β^n^*, the polynomial B-spline basis of degree *n* [15].

Using this representation, SplineDist predicts an object probability for each image pixel (*i,j*) along with a set of *M* two-dimensional control points **C**_*ij*_ [*k*] that fully define the spline curve (3). Similar to the StarDist radii, the **C**_ij_ [*k*] are expressed in relative coordinates with respect to (*i,j*), in polar form. More specifically, SplineDist predicts *M* angles 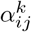 and *M* radial distances 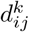 such that 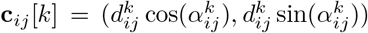 (Figure 1c). This construction is more flexible than star-convex polygons: in contrast to the *R* scalar radii at fixed angles, the *M* twodimensional control point vectors can point in any direction (angle and distance) from (*i, j*). The continuously-defined curve **s**_*ij*_ generated by the control point sequence {**C**_*ij*_ [*k*]}_*k*=0,... *M*-1_ can be sampled at any rate. We denote as 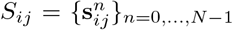 the set of *N* discrete points obtained by uniformly sampling s*_ij_* (*t*) as

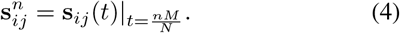

It is essential that we can construct a sequence *S* of any desired number of points *N* from {**c**_*ij*_ [*k*]}_k=0,...,M-1_ for two reasons. First, reference annotations come in the form of instance pixel masks, from which discrete contours can efficiently be extracted during training using classical image processing methods such as Satoshi and Abe’s algorithm [16].

Second, several control point sets may draw the same outline, so the very concept of ground truth control points does not make sense. We thus design SplineDist’s loss to evaluate the similarity between *S_ij_*, the discrete object outline that {**c**_ij_ [*K*]}_*K*=0,...,*M*-1_ generates, and a pixel-based ground truth object outline 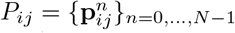 extracted from a reference instance mask. As a consequence, the value of *N* in (4) is dictated by the number of points in the ground truth *P_ij_*. In other words, SplineNet predicts the parameters of the continuous spline curve (3) that, when uniformly sampled, matches most closely the ground truth pixel outline of the object enclosing (*i,j*). SplineDist’s loss is formally expressed as

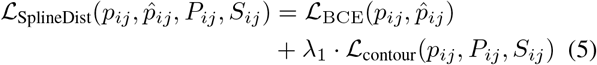

with

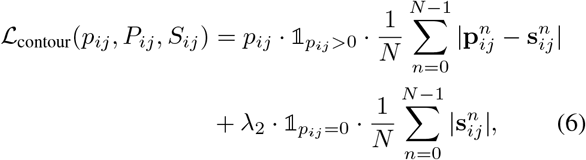

in which *p_ij_* and 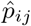 are the ground truth and predicted pixelwise object probabilities as in (1).

Because (5) involves the outline generated by the object model parameters and not the parameters themselves, a major difference with respect to StarDist is that individual pixels do not require to have their own ground truth. All pixels (*i, j*) enclosed in the same object 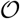 can indeed share a single ground truth outline 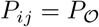 for all 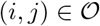 expressed in absolute (image) coordinates. The individual outline *S_ij_* that each pixel predicts can also straightforwardly be expressed in absolute coordinates by shifting the spline curve *s_ij_* around (*i, j*). This trick relieves the need to store relative coordinates describing the ground truth for each pixel. We can then afford considering the entire ground truth contour in the loss (as opposed to only *R* points) while remaining computationally tractable. The rest of SplineDist’s architecture is similar to StarDist, the remaining difference being the size of the final output layer (Figure 2).

**Fig. 2:**
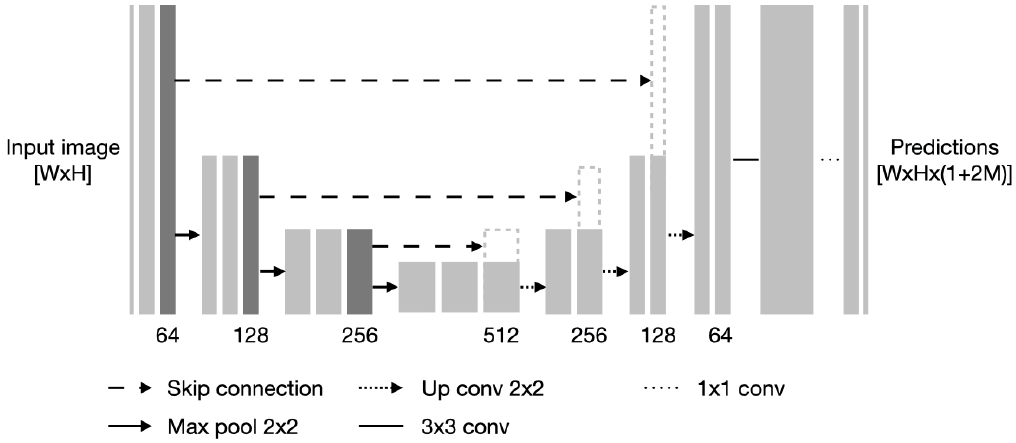
SplineDist architecture. For each image pixel (*i,j*), SplineDist densely predicts an object probability *p_ij_* and a sequence of *M* two-dimensional control points {**c**_*ij*_ [*K*]}_*k*=0,...,*M*-1_. The overall architecture mimics StarDist up to the dimensions of the output layer, sized *W* × *H* × (1 + *R*) in StarDist and *W* × *H* × (1 + 2*M*) in SplineDist, with *W* and *H* the input image width and height, respectively.

## 3. EXPERIMENTAL RESULTS

### 3.1. Implementation Details

We use the high-quality StarDist codebase (github.com/mpicbg-csbd/stardist) and mainly modify modules associated with ground truth generation and loss computations. For technical reasons, our implementation sets *N* as the number of points in the largest ground truth contour in the training set, and carries out subpixel interpolation to expand all shorter ground truth contours to this size. Hence, all pixels can be processed in a single tensor in the loss without unpacking, which critically speeds up training time. Like StarDist, SplineDist is implemented in TensorFlow [17] and is available at github.com/uhlmanngroup/splinedist.

### 3.2. Experiments

#### 3.2.1. Benchmarking

We compare the segmentation performance of SplineDist against StarDist on the image set BBBC038v1 (also known as Kaggle 2018 Data Science Bowl dataset), available from the Broad Bioimage Benchmark Collection [18]. It is composed of a diverse collection of images of cell nuclei which faithfully reflects the variability of object appearance and image types in bioimaging. BBBC038v1 is designed to challenge the generalization capabilities of a method across these variations and is therefore widely used for benchmarking. We employ the curated subset of BBBC038v1 provided in StarDist’s github repository, which is used in [7] to benchmark StarDist against state-of-the-art alternatives. This dataset is composed of 447 training images and 50 testing images.

Acknowledging the good approximation properties of cubic B-splines [19], we consider SplineDist_*β*^3^_ in our experiments. For a fair comparison, we use StarDist’s default network hyperparameters, data augmentation strategy, and partition ratio of training data into train and validation sets. We also used default training settings both in StarDist and SplineDist_*β*^3^_ (decaying learning rate of 0.0003, 400 epochs, batch size of 4). Finally, we use StarDist’s default NMS and object probability thresholds for both methods. We report the classical Intersection over Union (IoU) metric, also referred to as Jaccard index [20], to quantify instance segmentation quality.

In Figure 3, we quantitatively compare the performance of StarDist and SplineDist_*β*^3^_ with an increasing equal number of points (*R* = *M*). As expected from spline approximation theory [21], both methods eventually converge to the same results as the number of points grows larger since StarDist can be seen as SplineDist_*β*^1^_ with control points at fixed angles. The performance plateau, reached by StarDist from *R* = 16 onward, is already attained by SplineDist_*β*^3^_ at *M* = 6. The same segmentation quality can thus be obtained with 25% less output parameters (since SplineDist predicts 2*M* values for objects composed of *M* points). Increasing *M* further does not significantly improve performance. Because objects in BBBC038v1 tend to be small, large values of *M* result in spline curves with too many degrees of freedom, even causing a slight performance degradation for *M* > 12. In Figure 4, we show a visual example of results for *R* = *M* = 6. The smoothness granted by cubic B-spline approximation, coupled with the ability to predict the angular and the radial components of the model’s parameters, boosts SplineDist_*β*^3^_ performance for small *M* values.

**Fig. 3:**
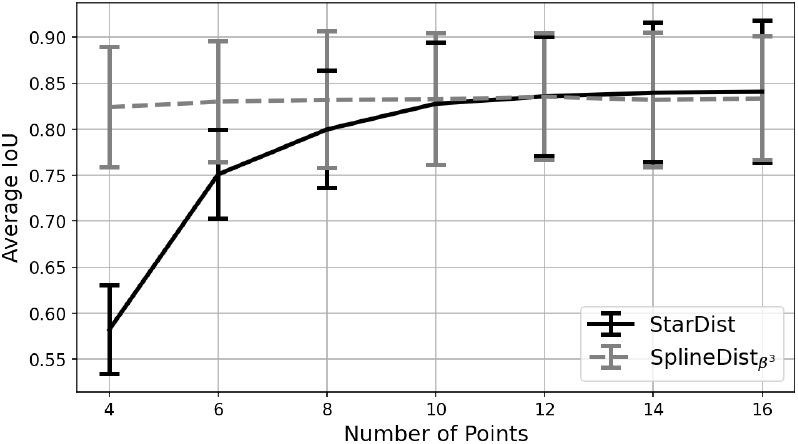
Evolution of the average IoU on the BBBC038v1 dataset for StarDist and SplineDist_*β*^3^_, with object models of increasing equal number of points (*R* = *M*). Error bars correspond to the IoU standard deviation on the whole dataset.

**Fig. 4:**
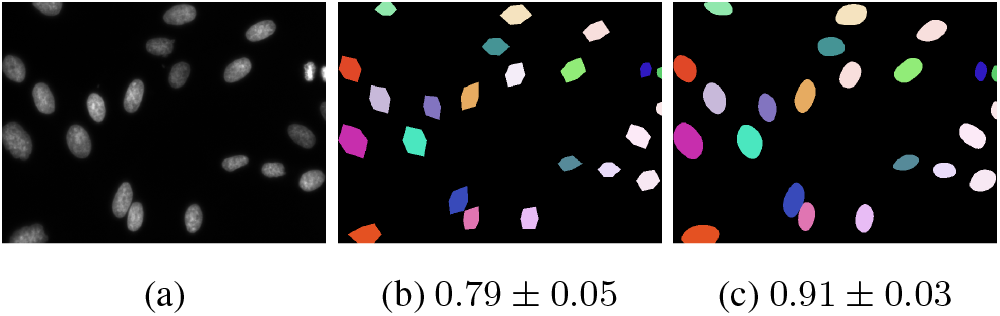
Visual comparison of instance segmentation results. (a) Original BBBC038v1 image, results obtained with (b) StarDist and (c) SplineDist_*β*^3^_ for *R* = *M* = 6. We indicate under each image its average IoU and standard deviation.

#### 3.2.2. Segmentation of non-star-convex objects

Because it can predict control points anywhere around the object and doesn’t rely on predefined angles, SplineDist can generate non-star-convex outlines. There is indeed no restriction imposing that all control points should have different angles. To explore this difference, we compare the performance of StarDist and SplineDist on a set of synthetic images containing mostly star-convex and some non-star convex cell-like objects. In that way, StarDist can still converge and we do not explicitly enforce SplineDist to learn how to represent non-star-convex objects. Our synthetic dataset is composed of 500 grayscale (8-bit) 512 × 512 images featuring from 1 to 20 randomly-shaped, non-overlapping deformed binary blobs that are subsequently degraded by non-uniform illumination, additive Poisson-Gaussian noise, and Gaussian blurring. We randomly split the dataset into 450 images for training and 50 for testing.

We compare the vanilla StarDist (*R* = 32) against SplineDist_*β*^1^_ with *M* = 32 (also a polygon of 32 vertices, but without the star-convex restriction) and SplineDist_*β*^3^_ with *M* = 8. The performance of SplineDist_*β*^1^_ reflects what could be expected from StarDist if radii angles were not fixed, while those obtained with SplineDist_*β*^3^_ illustrate the advantage brought by a smoother spline basis. While the average IoU of the three methods do not significantly differ since most objects in the dataset have pure or nearly star-convex shapes, a more in-depth assessment of the results (Figure 5) reveals a clear difference in performance on non-star-convex objects. StarDist either misses parts of non-star-convex objects or breaks them into several instances. We also note that SplineDist_*β*^3^_ achieves quantitatively equivalent and less noisy results than SplineDist_*β*^1^_ with 4× fewer points.

**Fig. 5:**
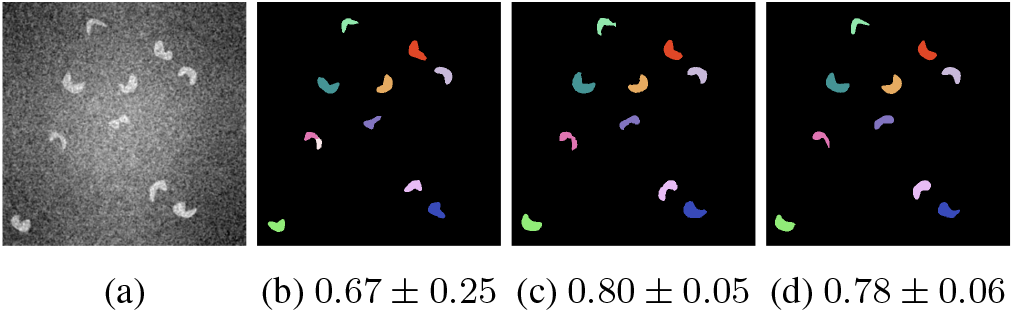
Segmentation of non-star-convex objects. (a) Original synthetic image, results obtained with (b) vanilla StarDist (*R* = 32), (c) SplineDist_*β*^1^_ (*M* = 32), and (d) SplineDist_*β*^1^_ (*M* = 8). We indicate under each image its average IoU and standard deviation.

## 4. CONCLUSIONS

We have introduced SplineDist, a modification of StarDist in which objects are represented as planar spline curves. SplineDist produces segmentation outlines of equally good quality as StarDist with an object model involving fewer parameters. Additionally, SplineDist generalises StarDist by lifting the star-convex polygon limitation. SplineDist therefore combines the detection performance of StarDist with the segmentation quality of spline curve models as traditionally used in parametric AC, while alleviating the weaknesses of either of these methods alone.

## Acknowledgements

The authors would like to thank Martin Weigert for inspiring discussions and helpful comments on this work, and Julien Fageot for valuable comments on the manuscript. This work is supported by EMBL core funding. The authors have no relevant financial or non-financial conflict of interest to disclose.

## Compliance with Ethical Standards

This is a computational study for which no ethical approval was required.

